# Fast and slow cortical high frequency oscillations for cortico-cortical and cortico-hippocampal network consolidation during NonREM sleep

**DOI:** 10.1101/765149

**Authors:** Adrian Aleman-Zapata, Richard GM Morris, Lisa Genzel

## Abstract

Memory reactivation during NonREM-ripples is thought to communicate new information to a systems-wide network. Cortical high frequency events have also been described that co-occur with ripples. Focusing on NonREM sleep after different behaviors, both hippocampal ripples and parietal high frequency oscillations were detected. A bimodal frequency distribution was observed in the parietal high frequency events, faster and slower, with increases in prefrontal directionality measured by Granger causality analysis specifically seen during the fast parietal oscillations. Furthermore, fast events activated prefrontal-parietal cortex whereas slow events activated hippocampal-parietal areas. Finally, there was a learning-induced increase in both number and size of fast high frequency events. These patterns were not seen after novelty exposure or foraging, but occurred after the learning of a new goal location in a maze. Disruption of either sleep or hippocampal ripples impaired long-term memory consistent with these having a role in memory consolidation.

## Introduction

Sleep is important for memory consolidation (*1*). Critical network interactions associated with systems consolidation are thought to occur during sleep, but only for salient information that needs to be abstracted across multiple events (*2, 3*). Reactivation of previous experiences during sleep is a long known phenomenon that has been proposed to enable systems consolidation (*4, 5*), but classically it is measured after performing on overtrained tasks. Such tasks, including track-running, may not necessarily induce post-training systems consolidation processes since no information needs to be extracted and the principle rule has already been learned. The distinction between new learning and overtrained tasks is critical as new research already calls many findings on memory reactivations into question when considering more behaviors involving relevant learning experiences (*6-8*). We retain memories of what is new and relevant to updating our model of the world. But what makes new information salient enough to trigger memory consolidation processes during sleep?

We recently proposed that dopamine coming from the locus coeruleus (LC) and ventral tegmental area (VTA) to the hippocampus could determine the fate of memories (*9*). Novel experiences sharing some commonalities with past ones (‘common novelty’) would activate the VTA and promote semantic memory formation via increased reactivations during sleep and ensuing systems consolidation (*2, 9*). By contrast, experiences that bear only a minimal relationship to past experiences (‘distinct novelty’) would activate the LC to trigger strong sleep-independent, initial memory consolidation in the hippocampus, resulting in vivid and long-lasting episodic memories (*2, 9, 10*).

Early ideas on memory were focused on multiple specialized areas being responsible for distinct forms of memory. More recently, we have moved on from thinking of one type of memory “residing” in a single brain area, to considering whole brain networks with dominant hubs in an encoding/retrieval network that determine the type or quality of a memory. In this framework, systems consolidation may induce a shift from an initial hippocampal hub to a primarily cortical one (*11*). The anatomical position of hippocampus allows it to orchestrate a wide range of cortical and subcortical networks and thus link various aspects of a given experience that are represented in distributed neocortical modules (*12*). This property would play a role during consolidation (and later reconsolidation) of the global memory network, but what be its physiological basis? An emerging idea is that hippocampal ripple oscillations during sleep are associated with memory reactivation across the whole brain and may be the potential mechanism for transmitting new information to whole-brain memory networks and adapting them (*13-15*). However, this new information would then subsequently need to be processed in the cortex.

The default-mode-network seems to have a special role within the global memory network, especially when considering semantic memories (*11*). The cortical areas within the default-mode-network connect the hippocampus to other cortical areas downstream such as the posterior parietal cortex that are critical targets for consolidation during ripples during sleep (*13, 16*). Thus consolidation would lead the dominant hub to shift along the default-mode-network: such as, for spatial memory, hippocampus to medial prefrontal cortex/retrosplenial cortex to downstream areas such as the posterior parietal cortex (*17-23*). Recent evidence showed that during hippocampal ripples other brain areas in the default-mode-network as well as the posterior parietal cortex also show high-frequency oscillations (*13*), which occur more often after learning. This global network synchronization would be the ideal state for global network adaptions with both increases and decreases of individual hub activity. Accordingly, ripple related activity has been shown to be able to both upregulate (*24*) and downregulate synaptic strength (*25*) on the cellular level. Thus, the ripple in the hippocampus as well as high-frequency oscillations in the cortex may be the functional cogwheels that enable the shift in the memory network and allow upregulation and adaption of cortical hubs and downregulation of hippocampal hubs during systems consolidation. Furthermore, cortical high-frequency oscillations may be the mechanism in the cortex for creating long-lasting memory representations.

Cortical interactions during and after hippocampal ripples are known to be critical for memory consolidation as seen both in the coherence across different oscillations (ripples, slow oscillations and spindles) (*26*) as well as co-occurrence of ripple events and high-frequency oscillations across brain areas (*13*). Unknown is how these network interactions change with a larger variety of behavior.

To test these ideas, we compared different behaviors in the event-arena (*27, 28*) to a non-learning “baseline” condition. (1) Foraging, a mix of open-field foraging and track running with small chocolate rewards spread along a track. This controls for the effect of food rewards, but contains no novelty. (2) Novelty, exploration of a new environment with novel cues/textures, a form of ‘distinct novelty’, which should lead to hippocampal cellular consolidation but no systems consolidation during sleep. (3) Plusmaze, training to a new reward location, which tests abstractions across multiple events (16 trials) in a familiar environment (with updated cues). Based on our hypothesis only the ‘common novelty’ of the Plusmaze condition should specifically lead to changes in post-behavioral sleep (*2, 9*).

## Results

We recorded electrophysiological signals for 4h in rats after four different conditions (Baseline, Foraging, Novelty, Plusmaze Fig. 1A). Each behavioral condition was performed in an event arena (1.5mX1.5m) (*27, 28*), which was adapted to each condition. For Foraging white curtains surrounded the arena and a divider was added to create a 1.5m track, which was 15 cm wide. Small chocolate cereal pieces were spread along the track to encourage the animal to move along it and control for effects of chocolate rewards. Once the animal had eaten the pieces the track was refilled so that the animal kept moving back and forth on the track over the 20min training period. For Novelty, the curtains were removed around the same event arena and it was filled with novel objects and textures such as bubble wrap and newspaper; the animal was allowed to explore freely for 20 min. For Plusmaze learning, the curtains were included and large cues were placed on the curtain, further the walls of the event arena were inverted so that a cross-shaped maze was created covering 1.5×1.5m with a track width of 15cm. First the animal could explore the Plusmaze freely for 10min with chocolate cereal rewards placed at the new goal. Then, the animal was trained for 15 trials from different starting locations and the goal arm baited with more rewards, which usually would take another 10min. For Baseline recordings no specific behavior was performed. The behavioral conditions (together with the later described conditions of SWR-D and Con-D) were counterbalanced across animals (see methods). After each behavior the animals were placed in a recording box and given a 4h sleep period during which electrophysiological signals were recorded. All animals had tetrodes targeting the dorsal hippocampus (AP-3.2, ML 2) and a screw electrode (ECoG) touching the posterior parietal cortex (PPC, AP -4.5, ML 5) as well as one above the prefrontal cortex (PFC, AP 3.5, ML 0.5) implanted. These signals were used to manual sleep score the 4h period (scorer blinded to condition, average ± SEM NonREM 93.34min ±3.45, Transitional 2.6min ±0.78, REM 7.9min ±1.38, see Fig. S1) and only the NonREM sleep periods were used in the subsequent analysis.

**Fig. 1.**
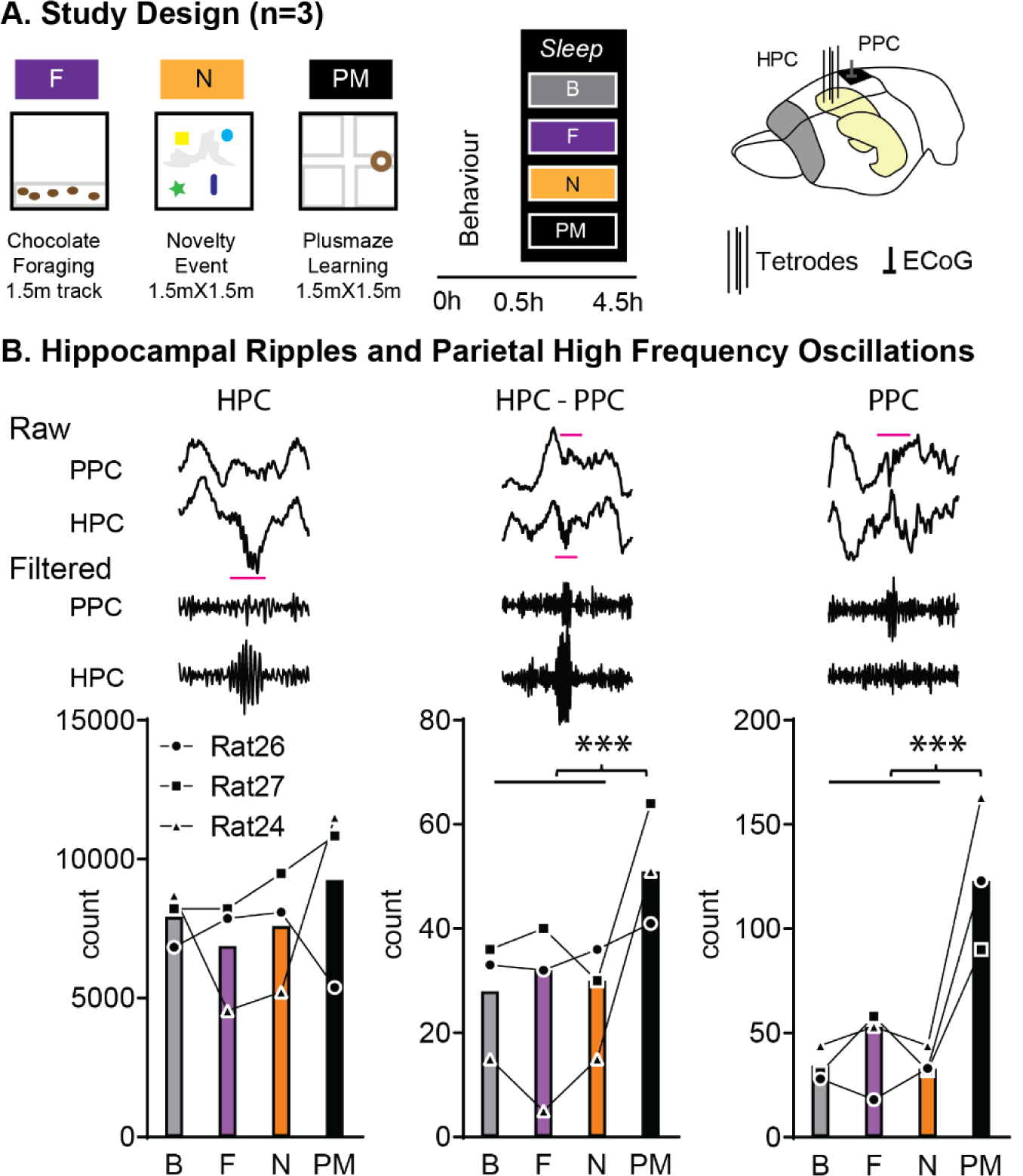
Behavior and NREM Ripples and High Frequency Oscillations. **A**. Study design. Three animals underwent three different behavioral conditions in the event arena: Foraging on a linear track with coco crumbs (F, 1.5mX1.5m open-field with novel objects/textures), and Plusmaze (PM, 1.5mX1.5m, 10min free exploration, then 16 trials to goal with chocolate cereal). We recorded a 4h sleep-period in after these behaviors and a non-learning Baseline (B). **B**. We detected ripple events in the hippocampus (AP-3.2, ML 2, tetrodes, yellow, **HPC**) and high frequency oscillations in the right posterior parietal cortex (AP -4.5, ML 5, ECoG, black, **PPC**) during NonREM sleep and classified events into single HPC, cooccurring HPC-PPC and single PPC. There were more HPC-PPC and more PPC events after Plusmaze than other behaviors. **Cond Effect** HPC p=0.66 F_3,6_=0.55, HPC-PPC p=0.037 F_3,6_=5.5, PPC p=0.003 F_3,6_=15.33, **Orthogonal comparisons:** p<0.001 Baseline (B), Foraging (F), Novelty (N), Plusmaze (PM).

### Ripples and high frequency oscillations

We focused on characteristics of NonREM-ripples in the hippocampus as well as posterior parietal cortex (PPC) high frequency oscillations (HFO) and detected these as discrete events. First, we divided them into three different type of events: individual hippocampal ripples (HPC), cooccurring hippocampal and cortical events (two events occurring within 50ms of each other, HPC-PPC) and single high-frequency oscillations in posterior parietal cortex (PPC). The occurrence of single hippocampal ripple events remained the same after all behaviors but after Plusmaze more HPC-PPC events and more single PPC HFO were detected than in the other conditions (Fig. 1B, condition: HPC p=0.66 F_3,6_=0.55, HPC-PPC p=0.037 F_3,6_=5.5, PPC p=0.003 F_3,6_=15.33).

Hippocampal ripples are known to occur as single events but also in groupings of doublets and more (multiplets). To check if it would be more likely to have HPC-PPC events during doublets and multiplets and if in general the occurrence of these were influenced by our different behavioral conditions, the hippocampal ripple events were classified as singlets, doublets and multiplets (triplets and more). Overall, HPC events were more likely occurring as hippocampal singlets (p=0.001 F_2,4_=84.69) but there was no difference for the occurrence of each type (singlets, doublets, multiplets) for HPC events across conditions (Fig. S2A). However, for HPC-PPC events in addition to being more prominent as singlets (p=0.014 F_2,4_=15.18) also showed a significant interaction between condition and type of hippocampal ripple (p=0.005 F_6,12_=5.62), with a condition effect only seen in HPC-PPC events if the hippocampal ripples were singlets but not in doublets or multiplets (p=0.013 F_3,6_=8.88, Fig. S2B).

To summarize, only cortical events showed a learning-specific increase and not individual hippocampal ripples or hippocampal ripple doublets or multiplets.

### Slow and fast high frequency oscillations

When investigating the PPC HFOs further, a bimodal frequency distribution was noticeable (Fig. 2A), which was not seen for hippocampal ripples. Thus next, we divided the PPC events by their individual theoretical determined split, which was at ∼155Hz for each animal. All types of events showed an increase after Plusmaze but the largest effect was seen in the fast (>155Hz) single PPC events (Full Model [HPC-PPC/PPC events, Slow/Fast, Cond]: cond Effect p=0.002 F_3,6_=17.94, HPC-PPC/PPC X Slow/Fast interaction p=0.036 F_1,2_=28.29, HPC-PPC/PPC X cond interaction p=0.019 F_3,6_=7.52 HPC-PPC/PPC X Slow/Fast X cond interaction p=0.039 F_3,6_=5.39 other p>0.19 F<3.6. Fig. 2B).

**Fig. 2.**
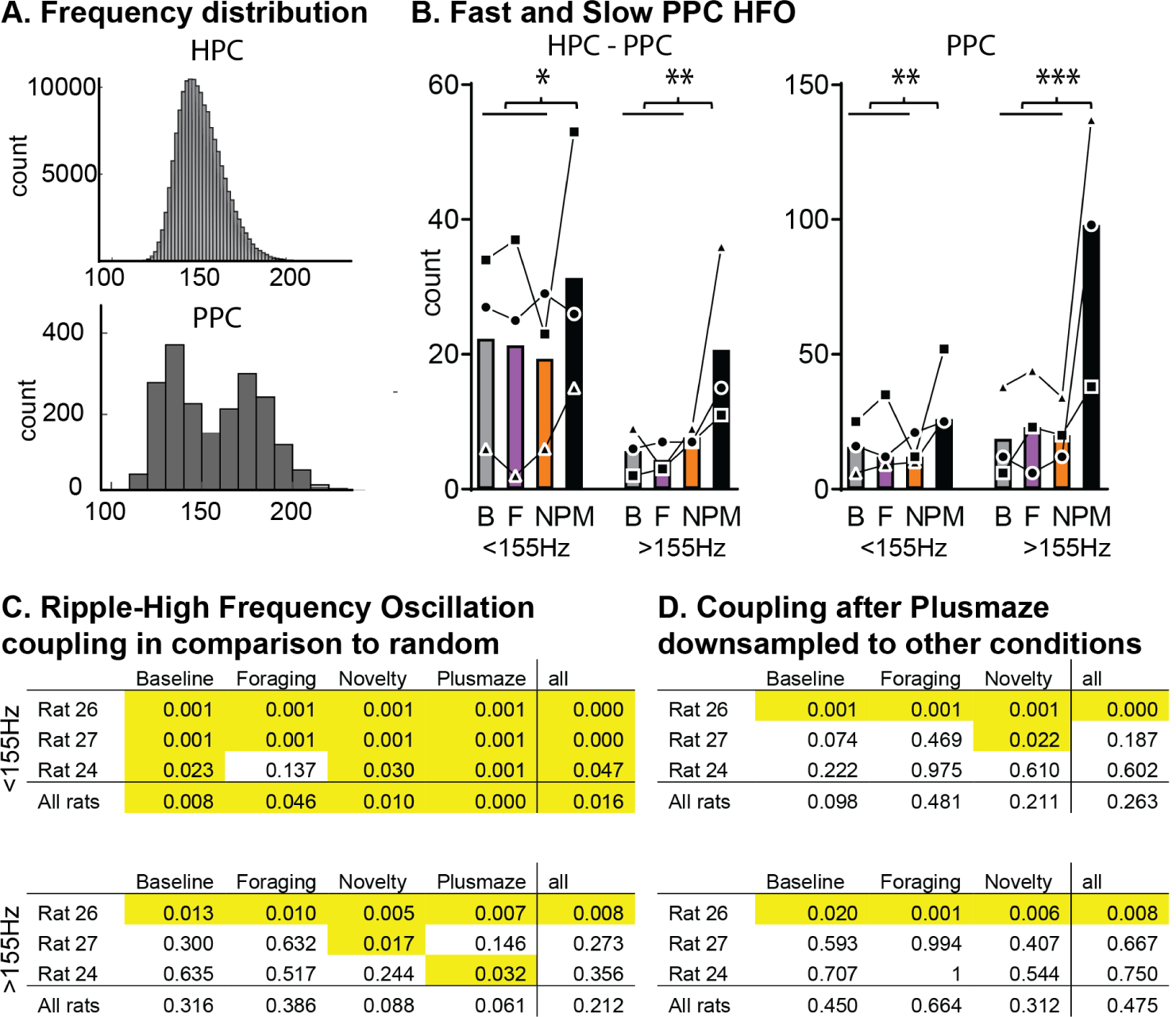
Fast and Slow High Frequency Oscillations. **A**. Histogram of hippocampal ripples (HPC) and parietal high frequency oscillations (PPC). For the latter the distribution was bimodal thus they were divided by ∼155Hz (individual threshold) and **B**. the largest increase after Plusmaze was seen in the fast single PPC events. **Full Model (HPC-PPC/PPC events, Slow/Fast, Cond)**: cond Effect p=0.002 F_3,6_=17.94, HPC-PPC/PPC X Slow/Fast interaction p=0.036 F_1,2_=28.29, HPC-PPC/PPC X cond interaction p=0.019 F_3,6_=7.52 HPC-PPC/PPC X Slow/Fast X cond interaction p=0.039 F_3,6_=5.39 other p>0.19 F<3.6. **Orthogonal comparisons**: *p<0.05, **p<0.01,***p<0.001. **C**. To check if cooccurrence of ripples and high frequency oscillations was above chance and not just random, we assigned random time-stamps to the high-frequency oscillations (per animal 1000 iterations) and checked for cooccurrence above this random distribution (p values in the tables, normalized within animal and condition, shown for each animal and condition separately as well as across condition and across animal). Only for the slower high frequency oscillations coupling to hippocampal ripples was above chance consistently. To check if the increase in coupling seen after Plusmaze was just caused by increase in count of events, we downsampled the number of events in the Plusmaze condition (1000 random iterations of downsampling) and compared the amount of coupling to each condition. Only one animal showed a higher rate of coupling after Plusmaze than in other conditions, thus increased coupling seen in B was mainly due to increase number of events less a learning effect per se. Baseline (B), Foraging (F), Novelty (N), Plusmaze (PM).

To check if the condition effect of HPC-PPC events and overall occurrence of HPC-PPC events was only a random biproduct of the occurrence of both HPC and PPC events, we performed two controls analysis. Firstly, we randomly assigned time-stamps within the NonREM periods to PPC events and checked in this simulated data for HPC-PPC events (1000 iterations) separately for slow and fast HFO. This simulation showed that cooccurrence of hippocampal ripples with slow PPC events was more than chance for each animal and condition, but this was not the case for the fast PPC events (Fig. 2C). Thus only the slow but not fast PPC events were significantly coupled to hippocampal ripples. Secondly, to check if the increase in HPC-PPC events after Plusmaze was just due to the overall increase in PPC events after Plusmaze or a specific, separate effect of learning, we randomly downsampled PPC events to the number occurring in each of the other conditions and then checked for cooccurrence. Surprisingly, this analysis showed that in two out of three animals there was no specific effect of Plusmaze on cooccurrences (both slow and fast PPC events), instead the increase after Plusmaze was only due to the general increase of PPC events after this condition (Fig. 2D).

To summarize, there seem to be two distinct types of PPC high frequency events: slower and faster ones. And especially single, fast PPC high frequency oscillations show an increase after learning. Further, only slow and not fast PPC high frequency oscillations were significantly coupled to HPC ripples and the increase of HPC-PPC events after Plusmaze was due to the overall increase of PPC events not a learning-effect in itself.

### Spectral profile of different events

The animals also had an ECoG place above the prefrontal cortex, but due to the anatomical distance between prelimbic, in which HFO have been shown to occur (*13*), and brain surface, individual HFO events could not be detected. To still be able to investigate any effects in the prefrontal cortex, we next switched to measuring spectral power in the same frequency range during the different types of detected events across all three brain regions. We split for single hippocampal ripples, as well as slow and fast PPC high frequency oscillations. Events classified as cooccurring PPC-HPC or single PPC did not show any differences, instead only differences was seen for slow versus fast PPC high frequency oscillations (see Fig. S3 for five event type split). For each type of event, the same number of events was included across all four behavioral conditions for each animal, the number included determined by the condition with the smallest numbers of events. Oscillatory power in both cortical ECoGs placed above the prefrontal and posterior parietal cortex as well as derived from the hippocampal tetrode targeting the ripple-range (100-250 Hz) (*13*) was extracted for ±50ms of event peak, as shown in Fig. 3. The power was normalized for each animal across all conditions and event types within each brain area to allow direct comparison of modulation across events. Overall the three event types differed in the spectral profiles across brain areas and there was a significant event type X brain area, type X condition and three-way interaction (Fig. 3, Full Model [BA, Type 3 levels, Cond]: type p=0.003 F_2,4_=31.93, BA p=0.007 F_2,4_=21.36, cond p=0.0856 F_3,6_=3.6, type X BA interaction p<0.001 F_4,8_=26.26, type X cond interaction p=0.035 F_62,12_=3.36, type X BA X cond p=0.058 F_12,24_=2.11). Ripples showed large hippocampal but less cortical power, slow PPC high frequency oscillations showed similar hippocampal power but more PPC cortical power in comparison to ripples. Finally fast PPC HFO showed larger PFC and PPC but smaller hippocampal spectral power (linear increase in power from ripple to slow and then fast PPC HFO for PPC p=0.007 and PFC p=0.005). The only condition effect was seen for fast PPC oscillations, there was an increase in power after Plusmaze in both cortical regions during these events (PPC cond p=0.011 F_3,6_=9.26, PFC cond p=0.034 F_3,6_=5.75).

**Fig. 3.**
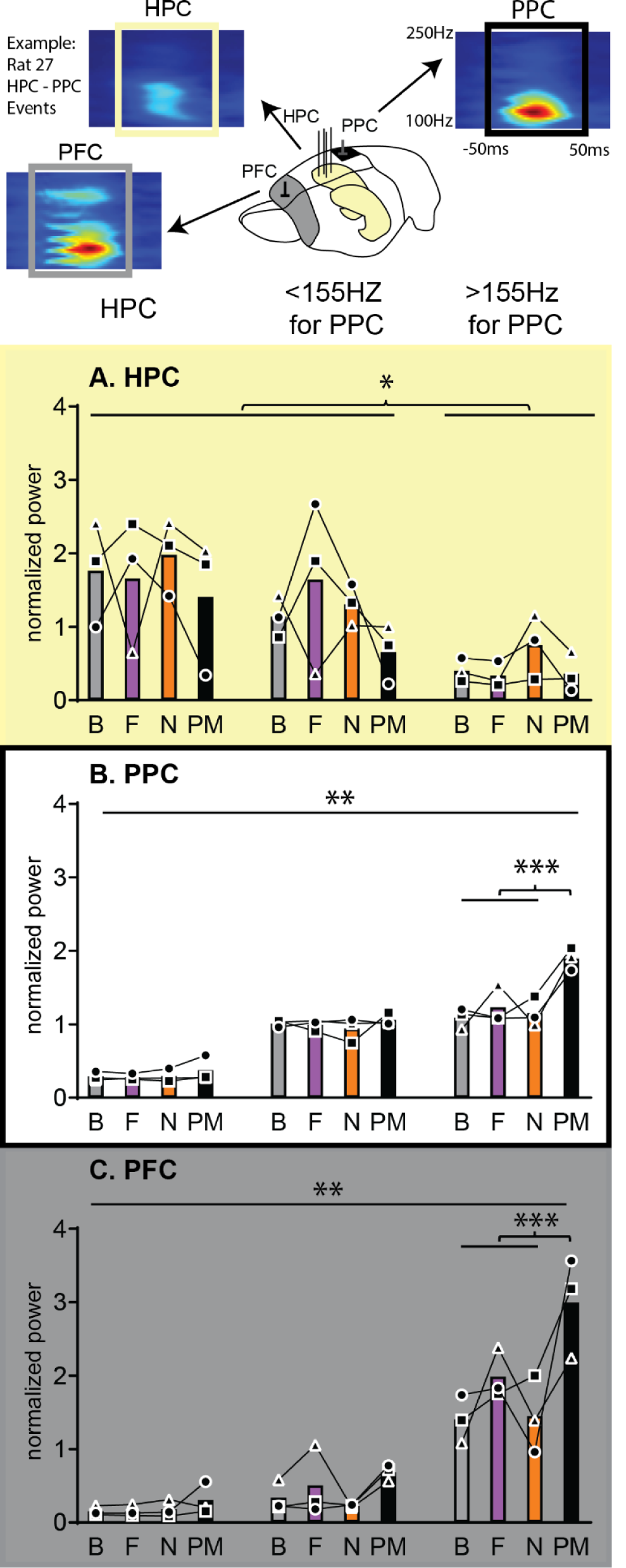
Spectral Power During Events: For all three brain areas normalized power. (for each animal across event type and brain area) for **100-250Hz ±50ms around the events** of the largest events in each condition (same number of events across conditions for each type). Shown for **A**. Hippocampus (HPC) **B**. posterior parietal cortex (PPC) and **C**. prefrontal cortex (AP 3.5, ML 0.5, ECoG, PFC). Overall the three event types differed in the spectral profiles across brain areas. Ripples showed large hippocampal power but less cortical, slow PPC high frequency oscillations showed similar hippocampal power but more cortical power than ripples. Finally fast PPC HFO showed larger PFC and PPC but smaller HPC spectral power. Cortical power both in PPC as well as PFC showed a linear increase from ripple to slow and then fast PPC high frequency oscillations. These general effects were the same if separated for those cortical events that were classified as coupled or not coupled to hippocampal ripples (see Fig. S3). As for condition, there was an increase in power after Plusmaze in PPC and PFC for the fast cortical events. **Full Model (BA, Type 3 levels, Cond)**: type p=0.003 F_2,4_=31.93, ba p=0.007 F_2,4_=21.36, cond p=0.0856 F_3,6_=3.6, type X BA interaction p<0.001 F_4,8_=26.26, type X cond interaction p=0.035 F_62,12_=3.36, type X BA X cond p=0.058 F_12,24_=2.11 (all other p>0.289). **for each brain area HPC:** type p=0.039 F_2,4_=8.10, **PPC**: type p<0.0014 F_2,4_=130.86, cond p=0.006 F_3,6_=11.98, typeXcond p=0.009 F_6,12_=5.00 **PFC:** type p<0.001 F_2,4_=1050.32, cond p=0.055 F_3,6_=4.53, typeXcond p=0.008 F_6,12_=5.18 **Just including fast PPC events**: PPC cond p=0.011 F_3,6_=9.26, PFC cond p=0.034 F_3,6_=5.75 **Orthogonal comparisons for condition**: *p<0.05, **p<0.01 Baseline (B), Foraging (F), Novelty (N), Plusmaze (PM).

In sum, fast PPC high-frequency oscillations, showed larger PFC and PPC and smaller hippocampal spectral power and a condition effect, with an additional increase in cortical power after Plusmaze. The effects in hippocampal spectral power across events corresponds well to the cooccurrence analysis (Fig.2), in which only slow but not fast PPC events showed above chance coupling to hippocampal ripples. Overall this analysis indicates that fast events correspond to a prefrontal-parietal network and slow events to a hippocampal-parietal network, and only the former and not the latter increase in size specifically with learning.

### Spindle analysis

Spindles are often reported to occur after hippocampal ripples (*26*) as well as ripples occurring in the throughs of the spindle (*29*). Further, cortical high frequency events have been reported to be associated with spindles (*13*). Finally, spindle number and size increase after learning events in humans (*30, 31*). From human subjects we also know, that one can divide spindles into frontal and parietal spindles, which each show different characteristics in individual typical occurrence rates and average frequency (*32*). Typically, after learning the faster, parietal spindles show increases, at least this has been shown in humans but not yet in rodents. Thus next, we detected spindle events in the PFC and PPC and could show that PPC but not PFC spindles increased after all behavioral conditions in contrast to baseline (Fig. 4A, brain area X cond p=0.02 F_3,6_=7.22). Coupling between ripples and both types of spindles (Fig. 4B) as well as PPC high frequency oscillations and PFC spindles (Fig. 4D) did not show any changes across conditions. However, coupling of both fast and slow high frequency PPC events and PPC spindles increased after Plusmaze and overall there were more coupling between fast events and PPC spindles (Fig. 4C, cond p=0.04 F_3,6_=5.29, type p=0.058 F_1,2_=15.61 interaction p=0.25). Since as with ripple-HFO coupling this could just be a result of a general increase in number of events after learning with both spindles and HFO showing such an increase, we performed a control analysis comparing cooccurrence in comparison to random time-stamps within the NonREM periods for the spindles as was done for ripple-HFO coupling. Surprisingly, for the high frequency events the coupling never was above chance, while for PPC spindle-hippocampal ripple coupling two out of three animals showed coupling, while the third animal only showed this effect after Novelty (Fig. 4F). The significant coupling between ripples and PPC spindles after Novelty was also seen for ripples occurring before spindles, but not for ripples occurring after spindles (Fig. 4G).

**Fig. 4.**
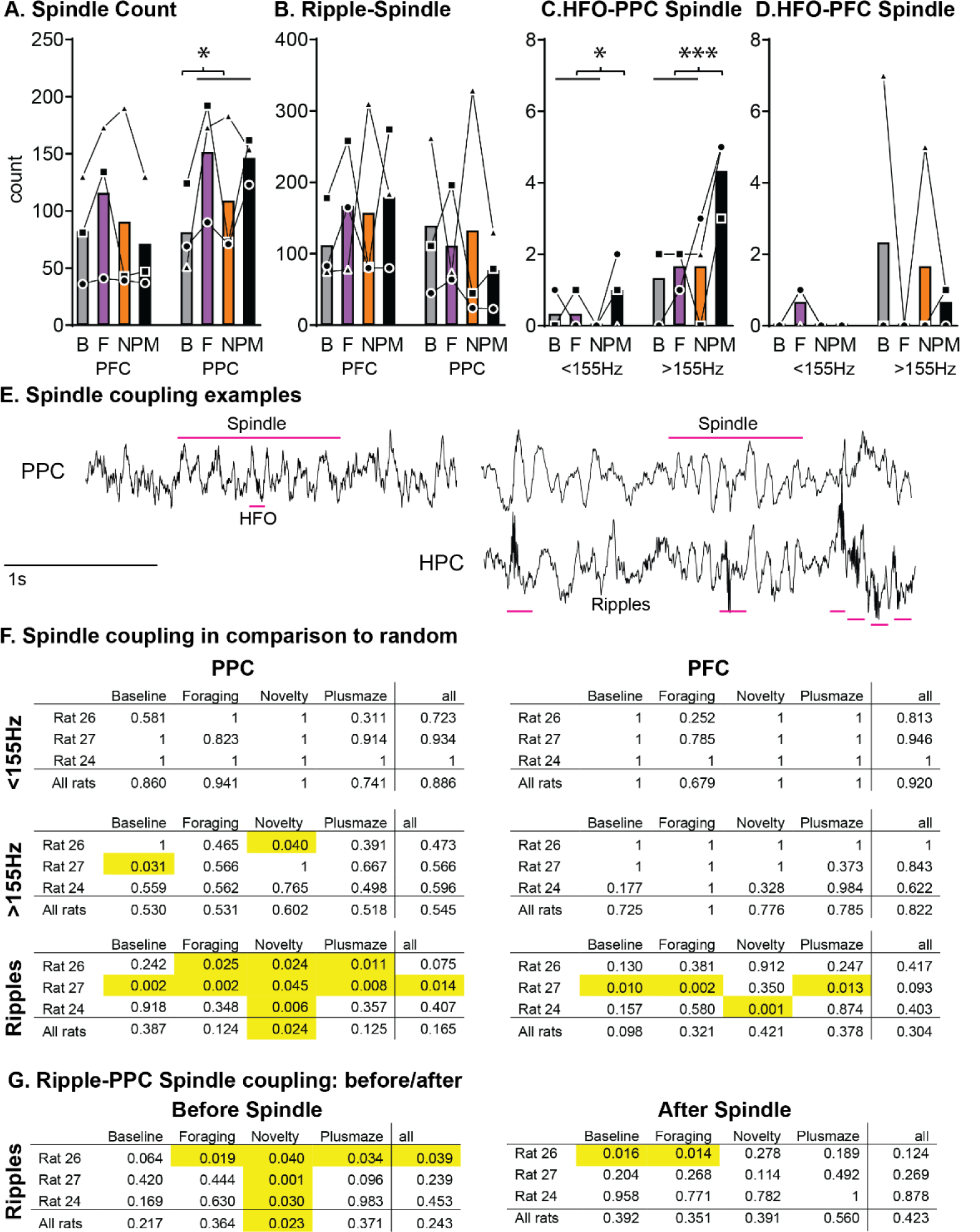
Spindle coupling During Events: **A**. We detected spindles in the two cortical areas (prefrontal cortex PFC and parietal cortex PPC). All three behaviors caused an increase in spindle events but only in PPC (brain area X cond p=0.02 F_3,6_=7.22, *p<0.05 orthogonal comparison). **B**. Across conditions there was no change in spindle-ripple coupling, but there was an increase in high-frequency oscillation to spindle coupling after Plusmaze for both slow and fast events but only for PPC spindles (**C**, cond p=0.04 F_3,6_=5.29, type p=0.058 F_1,2_=15.61 interaction p=0.25) and not for PFC spindles **(D). E**. Example traces for spindle coupling with high frequency oscillations and ripples. **F**. Shows the spindle to fast, slow HFO and ripple coupling (p-values) in comparison to random (1000 iterations with random time stamps for spindles) for each animal and condition as well as summarized over each. There was no significant coupling with the exception of PPC spindle-ripple coupling especially after Novelty. **G**. To test how specific this was, we took for each spindle the time before and after the spindle and checked for the same coupling. The coupling after Novelty could also be seen with ripples occurring before PPC spindles, but not for those occurring after spindles. For E and F: a p values of 1 means there were no cooccurrences. Baseline (B), Foraging (F), Novelty (N), Plusmaze (PM).

In sum, we could show that any type of behavior will lead to an increase in parietal but not prefrontal cortex spindles. Further, after Plusmaze there were more cortical high frequency oscillations occurring during parietal ripples, even though the coupling of these events was not above chance. Only ripples showed above chance coupling to parietal but not prefrontal cortex spindles, but also not reliably so. This coupling was present both for ripples occurring before as well as during spindles.

### Percentage of events

Until now, we used the absolute number of events in our cooccurring analysis. To enhance intuition how likely certain cooccurring events are, we next calculated percentages of these events in relation to the absolute number of each oscillation (Fig.5). This analysis is especially important when one considers how different the absolute number of events are with hippocampal ripples averaging across animals and conditions at 7907 events, slow HFO at 44 events, fast HFO at 49 events and spindles at 122 events during our sleep periods of 93 min. For PPC spindles 68% had a ripple occurring before and 69% a ripple during and 64% had a ripple after the spindle (pre/during/post p=0.025 F_2,4_=10.55, condition p=0.77 and interaction p=0.081). However, only each 2% of ripples occurred before, during or after PPC spindles. In regards to cortical HFO, only 0.35% of PPC spindles occurred with a slow HFO while 2% occurred with a fast HFO (slow vs fast p=0.068 F_1,2_=13.31, condition and interaction p>0.28); however 0.7% of slow HFOs and 7% of fast HFOs occurred during spindles (slow vs fast p=0.056 F_1,2_=16.48, condition and interaction p>0.8). Thus, while overall these events were very rare and not significantly different than chance, the fast HFO were more likely to be coupled to spindles than the slow despite the same amount of events for fast and slow HFOs. Finally, 51% of slow HFO were coupled to ripples but only 24% of the fast HFO (slow vs. fast p=0.031 F_1,2_=31.18, condition and interaction p>0.6). And only 0.35% of ripples were coupled to slow HFO and 0.11% of ripples to fast HFO (slow vs. fast, condition and interaction p>0.28).

In sum, most ripples were not coupled with either spindles or HFO, but a large percentage of spindles and HFO were associated with ripples. The difference is due to the vast difference in number of hippocampal ripples in comparison to the rare events of spindles and HFO (8000 vs 40-120). Thus one should imagine that the hippocampus just continuously shows ripples during NonREM sleep that then sometimes can cooccur with other events in the cortex. However, even after accounting for differences in absolute number of the individual events, it can be seen that ripples were slightly more likely to occur before and during PPC spindles than after, fast HFO were more likely than slow HFO to be associated to PPC spindles and slow HFO were more likely than fast HFO to be associated with ripples.

### Granger analysis of different events

Next, parametric Granger analysis on the same events types with a window size of 2.4s including all causality flows between hippocampus (HPC), prefrontal (PFC) and parietal (PPC) cortices was performed (Fig. 5; non-parametric Granger analysis see Fig. S4). We divided the analysis in two frequency ranges 0-20Hz and 20-300Hz. Slower oscillations are more specific for sleep and thought to represent coordination across brain areas and faster oscillation ranges information exchange. The event types did not show any differences across conditions in the granger analysis (Fig. S5); thus here we continued with averages across condition per animal. In the slower frequency ranges (0-20Hz) PFC→HPC and PFC→PAR showed an increase according to event types in granger values from HPC ripples to slower and then faster PPC events (Full Model with Freq range, Directionality, Event types: direct p<0.001 F_5,10_=24.85, types p=0.066 F_2,4_=5.76 direct X FR interaction p<0.001 F_5,10_=12.93, types X FR interaction p=0.0882 F_2,4_=43.75, direct X types interaction p<0.001 F_10,20_=12.79, direct X types X FR interaction p<0.001 F_10,20_=8.04 04 For each oscillatory band separately: 0-20Hz direct p=0.007 F_5,10_=6.28, types p=0.065 F_2,4_=4.43, direct X types interaction p<0.001 F_10,20_=11.111). In the faster frequency ranges (20-300Hz) PFC→PAR showed higher values for all types of events. And again PFC→HPC showed increases across the event types as already seen in the 0-20Hz frequency range (20-300Hz direct p<0.001 F_5,10_=54.97, types p=0.097 F_2,4_=5.83, direct X types interaction p<0.001 F_10,20_=7.71). As with the spectral power analysis, it did not make a difference if we divided the high frequency oscillation events into single or cooccurring (Fig. S4-6).

**Fig. 5.**
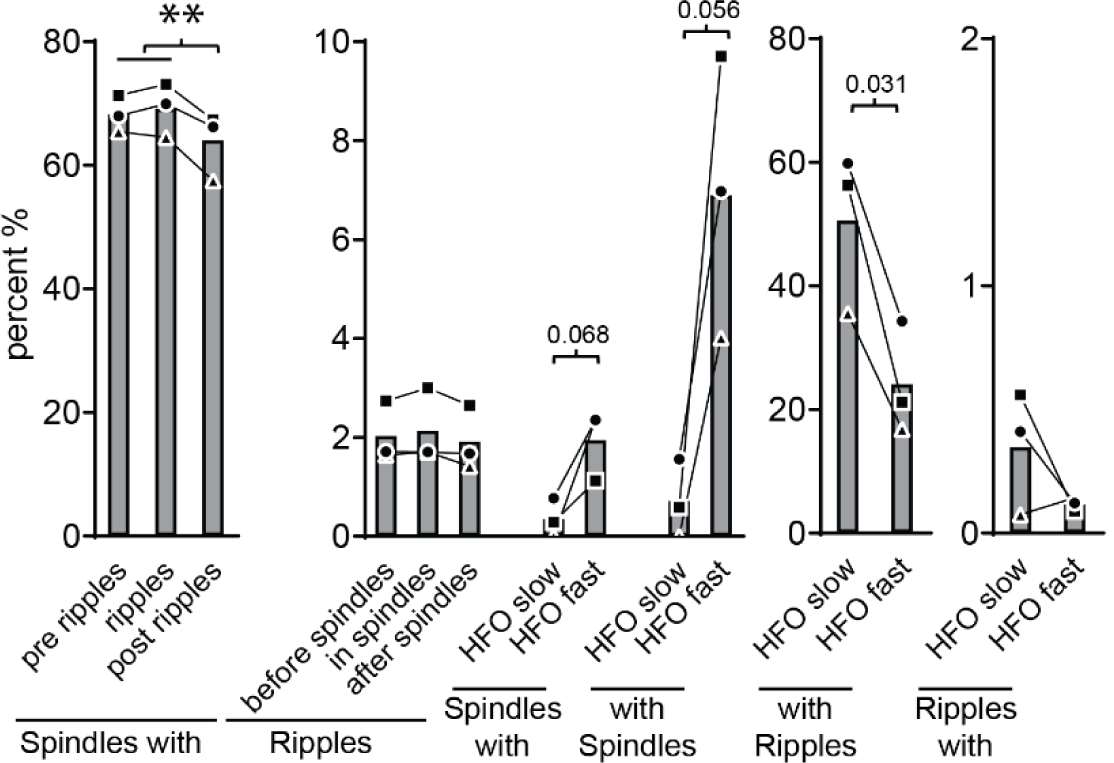
Spindle coupling Percentage of Events: We calculated the percentage of each cooccurring events, of note is the change in axis for the different event groupings. From left to right are the percent of PPC spindles that have a ripple before, during or after (**p<0.01 orthogonal comparison); percent of ripples that occur before, during or after a PPC spindle; percent of PPC spindles that have a HFO slow or fast oscillation (with p value); the percent of HFO oscillation that occur during PPC spindles (with p value); the percent of HFO that occur with a ripple (with p value); and the percent of ripples that occur with a HFO. These analyses are in relation to on an average of 7907 hippocampal ripples, 44 slow HFO, 49 fast HFO and 122 PPC spindles during our sleep periods of 93 min, statistical comparisons were only done for groupings with the same or similar number of events.

Thus, in sum especially faster PPC events showed a stronger lead of the PFC over other brain areas, which was independent if they occurred on their own or together with HPC ripples.

### Disruption by hippocampal ripple detection

The above analysis suggests that learning a new goal location in a Plusmaze changes prefrontal-parietal networks during NonREM high-frequency oscillation events and has less of an effect on hippocampal ripples. But is the activity of hippocampal ripples still necessary for long-term memory performance as seen in other experiments? To test this, we implanted animals with additional stimulating electrodes to the ventral hippocampal commissure (AP-1.3, ML 1, DV 3.8). Using similar methods others (*33*) have shown that disrupting hippocampal ripple activity daily (1h/d) slowed down learning in tasks trained over many days. Our one-session Plusmaze task allowed us to target a longer sleep period (4h) and compare this to a separate sleep deprivation group. Specifically, we compared 4h of sleep and sleep deprivation in unimplanted animals (within-subject, n=16, Fig. 6A) as well as sharp-wave-ripple-disruption (SWR-D), control-disruption (200ms after SWR, Con-D) and baseline (no stimulation, No-D) in implanted animals (within-subject, n=6, Fig. 6B). Animals performed above chance at 24h test (no food present) but performance fell to chance if sleep deprived after learning (Fig. 6A, cond p=0.052 F_1,15_=4.45, sleep to chance p<0.001, T_15_=4.56). SWR-D could mimic the sleep-deprivation effect, while both No-D and Con-D showed above chance performance (Fig. 6B, cond p=0.025 F_2,10_=5.42, to chance p=0.025 p=0.003). Thus sleep and more specifically activity related to NonREM-ripples is still necessary for long-term memory performance in this task.

**Fig. 6.**
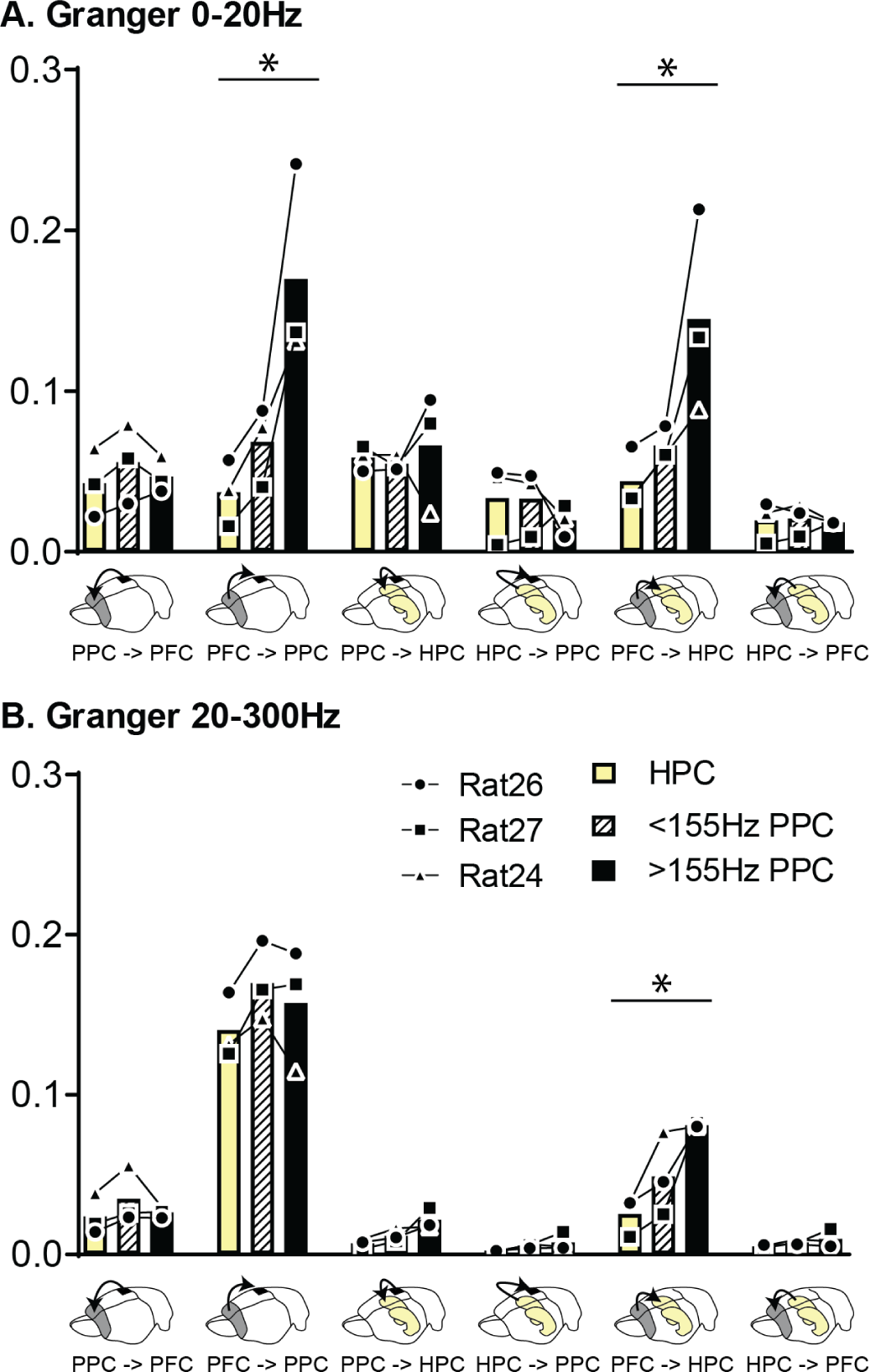
Granger Causality Analysis. (parametric) is shown for both 0-20Hz and 20-300Hz oscillation bands for the different event types (single hippocampal ripples HPC [yellow], slow [black and white shading] and fast [black shading] posterior parietal cortex high frequency oscillations PPC HFO) with the six possible directionalities. **A**. In the **slower frequencies** fast PPC HFO induced an increase in prefrontal cortex to hippocampal and to parietal granger values (linear increase from ripples, slow and fast HFO). **B**. In contrast in the **faster frequency** band overall PFC to PPC was increased for all events and PPC HFO showed an increase in PFC to HPC values (linear increase from ripples, slow and fast HFO). **Full Model (Freq range, Directionality, Events types)**: direct p<0.001 F_5,10_=24.85, types p=0.066 F_2,4_=5.76 direct X FR interaction p<0.001 F_5,10_=12.93, types X FR interaction p=0.0882 F_2,4_=43.75, direct X types interaction p<0.001 F_10,20_=12.79, direct X types X FR interaction p<0.001 F_10,20_=8.04 For each oscillatory band separately: **0-20Hz** direct p=0.007 F_5,10_=6.28, types p=0.065 F_2,4_=4.43, direct X types interaction p<0.001 F_10,20_=11.11, **20-300Hz** direct p<0.001 F_5,10_=54.97, types p=0.097 F_2,4_=5.83, direct X types interaction p<0.001 F_10,20_=7.71, **Type effect linear contrast**: *p<0.05. prefrontal cortex PFC, hippocampus HPC, posterior parietal cortex PPC.

**Fig. 7.**
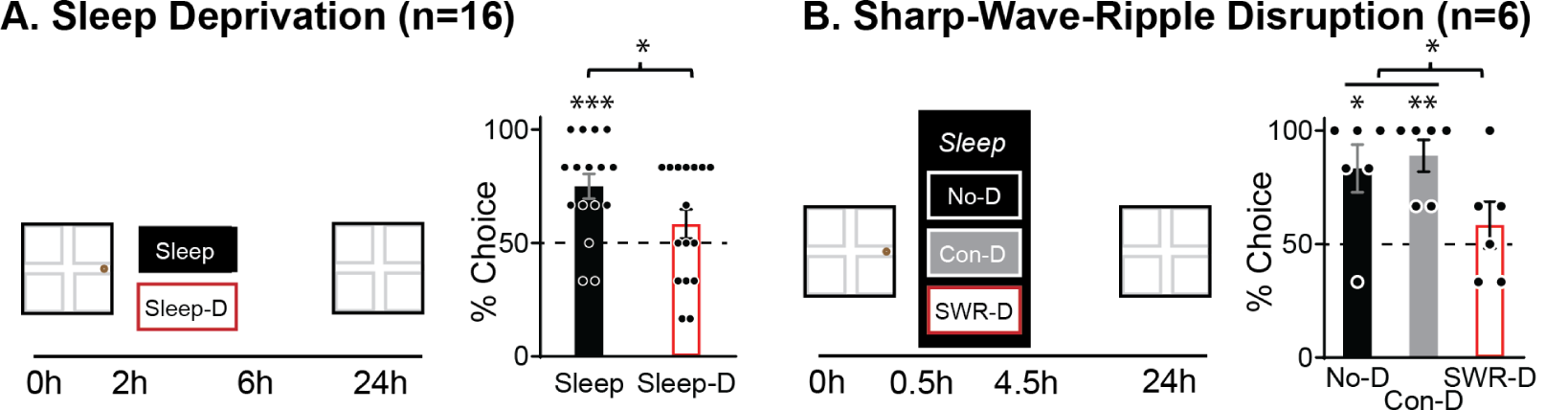
Sleep Deprivation and Ripple Disruption. **A**. Animals were trained in the Plusmaze and were either sleep deprived (gentle handling) or allowed to sleep for 4h and then retested 24h later (no food present). Only after sleep and not sleep deprivation (Sleep-D) could the animals remember the previous day’s goal location. cond p=0.052 F_1,15_=4.45, ***sleep to chance p<0.001, T_15_=4.56 **B**. As above but now implanted animals were trained in the Plusmaze and then received sharp-wave-ripple disruption (SWR-D), control-disruption (200ms delay, Con-D) or no-disruption (No-D) for 4h. Only with intact hippocampal ripples could the animals remember the previous day’s goal location. cond p=0.025 F_2,10_=5.42, to chance *p=0.025 ** p=0.003

The same ripple dependent, control disruption and sleep deprivation was performed after a memory-competition paradigm in the watermaze (*10*). In the watermaze dwell times at 24h test were not affected, thus it seemed that memory for the platform location remained intact across interventions. However, sleep deprivation as well as ripple-disruption did affect to which platform location the animals swam to first (Fig. S7). This indicates, that while in contrast to the Plusmaze task the watermaze memory was strong enough to survive ripple disruption and sleep deprivation, the memory representation was affected in comparison to the control and created a weaker behavior response.

## Discussion

The key observation is that learning – encoding a new goal location across multiple trials – induces changes across the hippocampal-prefrontal-parietal network during ripples and high-frequency oscillations in NonREM sleep, which was not seen after other behaviors such as experiencing novelty or foraging for chocolate. More specifically, learning led to an increase in high-frequency oscillations in the cortex. Two different types of cortical high-frequency oscillations could be identified: low and high frequency, which we categorized as below or above 155Hz. The faster high-frequency oscillations showed a learning-specific increase, occurring more often and becoming larger after Plusmaze learning as measured quantitatively using spectral power.

Further observations were that the fast high-frequency oscillations displaying higher cortical power (prefrontal and parietal) showed lower hippocampal power. In contrast, slower high-frequency oscillations showed more hippocampal and less prefrontal cortex power, and only for these slower events was the coupling to the hippocampal ripples above chance. Parietal but not prefrontal cortex spindles increased after all behavioral conditions, but only after Plusmaze learning did more high frequency events occur during parietal spindles. However, spindle-high frequency oscillation coupling was not above chance and was just a biproduct of more events after each behavior. In granger analysis the fast, cortical high-frequency events also showed increased lead of the prefrontal cortex over both the parietal cortex and hippocampus. Finally, while there was no learning-specific effect for the number or size of hippocampal ripples, they were still necessary for memory performance the next day. Both ripple-disruption and sleep deprivation led to memory falling to chance at test.

The seminal paper by Khodagholy et al (*13*) was the first to describe cortical high-frequency events. They observed these events in default-mode network regions (prefrontal and retrosplenial cortex) as well as the posterior parietal cortex, and these events tended to occur together with the hippocampal ripples. It was also seen that these cooccurring events became more common after learning. We replicated this observation in our learning paradigm, but now add that there seem to be two different types of cortical events – slower as well as the faster ones. These two different types could already be seen in the coherence figures for retrosplenial to parietal cortex in Khodagholy et al’s (2017) paper but not explicitly described in the article. Our data suggest that the faster events represent more of a prefrontal-parietal network with less hippocampal involvement and they are likely to be initiated by, or contain information from, the prefrontal cortex as seen in granger causality analysis. In contrast, the slower events represented parietal-hippocampal network interactions with less prefrontal cortex involvement. Further, learning a new goal location in a Plusmaze increases especially the faster cortico-cortical events in number and size.

Until now most memory research in rodents is focused on the hippocampus and amygdala, most likely due to the preferred usage of simple tasks (such as spatial learning and fear conditioning). Once more complex learning is involved the importance of the hippocampal long-term storage decreases, even if it is involved during initial learning, and instead memory storage relies increasingly on cortical areas (*28, 34*). Amongst cortical areas, the prefrontal cortex seems to take a special role when updating memories (*35, 36*) as well as coordinating consolidation during sleep across other cortical areas (*37*). Here, we could show that the prefrontal cortex leads the parietal cortex and hippocampus during the fast cortical high-frequency events that increase with learning. In contrast, the importance of the posterior parietal cortex during consolidation in our Plusmaze task, is most likely due to its spatial nature. The posterior parietal cortex serves as a cortical integration site for hippocampally generated allocentric spatial information and egocentric spatial orientation to permit goal-directed navigation (*13, 17-19*). Further, memory reactivation during sleep has been observed in the parietal cortex as in the prefrontal cortex (*15, 20*).

The main effect of learning a new goal location was seen in cortical high-frequency oscillations that occurred independently of hippocampal ripples. However, we also showed the necessity for the hippocampal ripples for memory performance during next day testing in our disruption experiment. It is thought that new information is transmitted to the cortex during hippocampal ripples that then has to be processed in the cortex to create a long-lasting memory trace. Perhaps it is this processing that is occurring in the cortical events. However, while these events would then show stronger learning related responses, the hippocampal ripples with their information content would still be necessary. It has been proposed that the hippocampus would record everything we experience throughout the day (*38*) and then during the night when we sleep our brains would sort through all these memories by reactivating them during hippocampal ripples but only postprocess and retain those memories that are recognized to be salient (*2*). This theory would fit to our results: hippocampal ripples remained the same after all experiences as one would expect if all that we do during the day is recorded, with reactivation largely restricted to sleep. But only after Plusmaze learning – a salient experience – did a significant amount of postprocessing of this new information occur during cortical high-frequency events.

We also investigated the association of these events with sleep spindles. Spindles are often reported to occur after hippocampal ripples (*26*) as well as have ripples occurring in their troughs (*29*). Further, cortical high frequency events have been associated with spindles (*13*) and spindles increase in number and size after learning events, at least in humans (*30, 31*). From humans subjects we also know, that one can divide spindles into frontal and parietal spindles, which each showing different characteristics in their individual typical occurrence rates and average frequency (*32*). After learning, the faster, parietal spindles are typically shown to increase. In contrast, most rodent studies only record from one site, and until now have tended to focus on frontal recording sites. Here we confirmed the increase of the parietal but not frontal spindles after behavior, but this was not specific to Plusmaze learning as it was was also seen after Foraging and Novelty. But only after Plusmaze learning was there an increase of parietal spindle and high frequency oscillation coupling. Interestingly this coupling was not above chance but only a biproduct of the increased number of events after Plusmaze, which was also seen in the ripple and high frequency oscillation coupling. Of course, that does not preclude that these couplings may still serve a purpose during memory consolidation, only that there seems to be no additional regulation increasing coupling after learning. It does highlight that one should be careful when discussing coupling between events that occur very often. We could confirm significant ripple-spindle coupling, which has often been reported, even though only two out of three animals showed this coupling to be above chance and it was mainly seen after Novelty not Plusmaze learning. Interestingly, coupling was significant both for ripples occurring before as well as during spindles in the Novelty condition. Many rodent researchers focus on the ripples before the spindle (*26*), while others especially those working with human intracranial data focus on ripples occurring during spindles (*39*). Here we could confirm both associations and ripples tend to occur before as well as during spindles, even though overall coupling was quite weak. Examples of these trains of events were already depicted in Maingret et al (2016) in which the artificial enhancement of the coupling of ripples to slow oscillations and spindles was shown after triggering stimulations of ripple events. Thus that study focused on the spindles occurring after the ripples, additional ripples could be seen occurring during the spindle. It would be tempting to speculate that the sequence of ‘ripple-before’ spindle and ‘ripple-within’ spindle events represents the dialogue between hippocampus and cortex, with the hippocampus initiating and “preparing” the cortex with the first ripple to receive another input during a spindle (next ripple). Here, this coupling was generally present, stronger after Novelty and not specific to learning events. Further, the association was not very reliably present across animals. This does fit to the fact that these coupling events are very rare, ∼100 ripple-spindle events (with ∼70% of PPC spindles being associated with ripples but only 6% of ripples occurring before, during or after PPC spindles) and ∼6 high frequency oscillation-spindle events (with 2% of spindles associated with fast HFO and 7% of the HFO associated with spindles) across an average of 90min of sleep. Also others report them as very rare events (∼2-10% of ripples are followed by spindles in (*26*)). Thus perhaps while interesting to investigate, they may have less importance for memory consolidation as often proposed or at least ripples must serve many more functions than just during this coupling.

Overall, these result highlight that it is critical to consider which behaviors are used to measure “learning” and post-learning sleep processes. Up until now in rodent research this term is loosely used to describe any of the behaviors harnessed in these experiments. Especially in electrophysiological experiments, simple tasks that can be repeated daily and result in many perfect performance trials are often preferred for practical reasons. However, our results emphasize that consolidation signatures in sleep can mainly be seen in tasks that correspond to significant learning i.e. the extraction of salient, novel information across multiple trials.

In sum, we could show that ‘common novelty’ – extracting a new goal location in a familiar maze across multiple trials – induces changes across the hippocampal-prefrontal-parietal network during NonREM ripples and high-frequency events. A different pattern was seen, in contrast, during ‘distinct novelty’ or very familiar behaviors (Baseline, Foraging). The learning effect was mainly expressed in fast cortico-cortical high-frequency events that increased in number as well as size. This is the first evidence supporting the occurrence of two distinct types of cortical high-frequency oscillation that represent different consolidation networks: fast high frequency events for cortico-cortical network and slow events for hippocampal-cortical. Finally, ripple activity and sleep after learning in the Plusmaze is necessary for long-term memory performance in this task. These results are also the first direct experimental support for the hypothesis that different types of novelty affect sleep related consolidation differently (*2, 3, 9*). Reactivations during sleep-ripples are thought to allow memory abstraction across multiple events, such as multiple trials or sessions in a learning task, and thus the consolidation from initial hippocampal to long-term cortical memory storage when we encounter something new that fits into what we know (*5*). In contrast, novelty events or a foraging tasks – as commonly used for reactivation analysis – due not elicit such a brain-wide response.

## Acknowledgments

This work was funded by a Branco Weiss Fellowship – Society in Science to Lisa Genzel, we would like to thank Francesco Battaglia, Federico Stella and Gio Piantoni for advice on the analysis and Francesco Battaglia for supplying the SWR-D script.

## Contributions

AAZ performed the data analysis, RGMM helped design the study and provided the lab-space, LG designed, performed and analyzed the experiment and wrote the first draft of the manuscript. All authors revised the manuscript.

## Methods and Materials

### Subjects

Lister-hooded rats of 2 month of age (Charles River) were group housed with a 12 h:12 h light:dark cycle and water and food were provided ad libitum. After surgery, animals were individually housed with environmental enrichment (wood blocks). Previous to the surgery, animals were handled and accommodated to eat ‘Weetos’ cereals (Kellog’s) for 5 consecutive days. The last three days of handling, rats were habituated to the sleep boxes (75 cm x 35 cm x 50 cm) containing bedding material for 8 h. Under isoflurane anesthesia, a 0.5mm AP x 0.5mm ML craniotomies in the right hemisphere were made above CA1 (AP: -3.2 mm, ML: 2 mm from bregma) for later placement of the tetrode drive (7 tetrodes, individually movable). One small screw (M1×4) was driven into the bone above the cerebellum as a ground electrode for recordings and other 4 additional screws were fixed to the skull to stabilize the structure. Two more screws (M1×4) were soldered to wires for ECOGs and implanted above right parietal cortex (AP -4.5, ML 5, PPC) and above the right prefrontal cortex (AP 3.5, ML 0.5, ECoG, PFC).

After carefully removing the dura mater, polyamide tubes bundles were placed on the cortical surface and subsequently the cortical surface was covered with sterile vaseline. Screws and microdive were cemented to the skull using quick adhesive cement (C&B Metabond®) and dental acrylic cement. In the next days to implantation, animals were placed in the sleep box, signal was checked and electrodes were gradually driven into the brain target area over the course of up to 1.5 weeks. After 2 weeks of recovery, rats were handled and habituated to the arena according to the protocol previously described. Once the target area was reached, the recording sessions started for all the conditions. Using the Open Ephys recording system electrophysiological recording took place for 4 h after training in the sleep box. The headstage plugging prior to recordings, did not cause any stress on animals due to the well handling and ‘Weeto cereals’ animal fondness.

### Behavioral training

#### Habituation

Animals were thoroughly handled in their second week after arrival in the animal facility. Each animal was actively handled daily for at least 5 minutes. We emphasize here that handling of the animals is extremely important. Next animals were habituated to the test apparatus: 3 days track running and multiple sessions in the sleep recoding box (each +2h). The sequence of the different conditions in the Plusmaze (SWR-D, Con-D, Baseline) were counterbalanced across animals and followed by Foraging, Baseline (home cage) and Novelty. The same was done for Sleep and Sleep deprivation in the unimplanted animals. If a condition was performed twice (e.g. No-D or Sleep/Sleep deprivation) the average of both performances was used for analysis.

#### Training

##### Plusmaze

Each session had a new goal arm and new cues on the curtain walls surrounding the maze. The first trial had 1.5 wheetos (chocolate cereal) at the end of the goal arm and animals were allowed to explore freely for 10 min, this was followed by 15 trials with 0.5 wheetos at the goal location with different starting locations. After the training animals were placed in the recording box (implanted animals) or sleep box/home cage for sleep deprivation (unimplanted animals) for 4h. In implanted animals the undisrupted, swr-d and con-d were run counterbalanced in sequence across animals, followed by the non-learning baseline, forage and novelty conditions. Sessions were either in the morning or afternoon, but time of day was kept constant for each animal. In unimplanted animals sleep and sleep deprivation was counterbalanced over animals and sessions (2 rounds of each per animal).

##### Novelty

Animals were placed in a 1.5mX1.5m event arena filled with novel objects, textures and smells and could explore freely for 30min and then were placed in the recording box for 4h.

##### Foraging

Animals ran along a 1.5m track with chocolate sprinkles and then were placed in the recording box for 4h.

### Data Analysis

#### Signal acquisition

The local field potential (LFP) signals of the hippocampus were recorded using bundles of four electrodes (tetrodes) while the animals were sleeping. Simultaneously, the intracranial electrocortical signals (ECoG) of the parietal lobe and medial prefrontal cortex were recorded using screw electrodes. The recordings used were monopolar to avoid the influence of an active reference. The acquisition and recording of the electrophysiological signals were performed using the Open Ephys multichannel acquisition board (*40*) which makes use of *Intan Technologies* RHD-series chips with a 16-bit analog-to-digital converter and a sampling frequency of 20 kHz.

#### Data pre-processing

All signals were downsampled to 1 kHz by making use of an anti-aliasing 3rd order zero-phase Butterworth low-pass filter and a decimation by a factor of 20. Furthermore, an exploratory analysis of the frequency spectrum of the brain signals showed power line artifacts around 50 Hz and its harmonics, which were filtered out using spectral interpolation with their neighboring frequencies as suggested by Mewett et.al. (*41*). A crucial step in this analysis was detecting the times when the ripples and cortical high frequency events occurred, which are typically found during the NonREM sleeping stage. With this aim, the recordings were manually inspected and blindly scored into four stages in 1 second epochs: Wake, NonREM, a transitional sleep period between NonREM and REM; and REM itself. The cumulative percentage of time spent in each stage for all behavioural conditions is shown in SFig. 1.

#### Event detection

Once the sleep stages periods had been identified, only those scored as the NonREM stage were extracted from the original recordings of all brain areas. In order to identify the ripples, the hippocampal recordings were bandpass-filtered on the ripple spectrum (100-300 Hz) with a 3rd-order zero-phased Butterworth filter. The resulting signals were used to find the ripples starting, ending and peak times by thresholding voltage peaks which lasted a minimum duration of 30 msec. The thresholds were determined following a visual inspection of the detections and for each animal the same threshold was used across conditions. Two detected ripple peaks closer than 50ms were considered a single ripple. The same procedure was done with the parietal recording to detect HFOs and a maximum duration criteria of 100 msec was included to control for microaurousals. HFOs were classified as slow or fast based on their mean frequency which was computed with the *meanfreq* Matlab function. The cutoff frequency used to classify the events was obtained by first fitting all events mean frequencies into a Gaussian mixture model with two components and a shared covariance using the *fitgmdist* function from Matlab. In addition, the mixture model was used to cluster the mean frequencies into two groups. The maximum mean frequency value found in the low frequencies cluster was identified as the cutoff frequency. This cutoff frequency was first computed per condition to later find a single mean value per rat which was used to split the events among all conditions. A comparison among rats resulted in a mean value close to 155Hz.

Spindle events were detected in PPC and PFC single channels by using an adaptation of the YASA sleep analysis toolbox spindle detection script (DOI: 10.5281/zenodo.2370600), which is inspired by the A7 algorithm from (*42*). The broadband and sigma frequency ranges were defined as (1-30 Hz) and (9-20 Hz) respectively. A relative power threshold of 0.3 was used for all channels while the thresholds for moving correlation and moving RMS were adapted for every rat and brain region following a visual inspection. For each rat the same threshold were used across conditions. Events which were above these thresholds were considered a spindle if they had a duration between 0.5 and 2 seconds. Spindles that were closer to 500 msec were merged together.

#### Cooccurrence of events

Cooccurrence was determined for ripples and high-frequency oscillations if their respective peaks were within 50 msec. Spindles cooccurring with ripples or high-frequency oscillations were detected if the timestamps of their respective durations overlapped. To check if cooccurrence was above chance, we compared to random simulation controls. For cooccurrence between HFOs and ripples the timestamps within the NonREM epochs were randomly permuted and the original sample corresponding to the HFOs peak was assigned with a new random timestamp. Cooccurrence was then computed and stored. These steps were iterated 1000 times to create a null distribution. The values of the distribution were then normalized per animal and condition by subtracting the original cooccurrence value and dividing by the standard deviation of the distribution. This resulted in distributions in which zero represented the cooccurrence value with the original timestamps. The p-value for each distribution was computed by finding the ratio between the amount of values equal or above zero and the total amount of values in the distribution. Random simulations of spindles cooccurring with ripples and HFOs were performed in a similar way as HFOs with ripples with the difference that the sample corresponding to the spindle start was assigned with a random timestamp on every iteration and the end of the spindle was determined by adding the spindle duration. Cooccurrence, normalization and the computation of p-values were performed as mentioned above. To determine the amount of ripples occurring before and after a spindle a period of the same duration as the spindle was considered before its start and after its end respectively.

#### Spectral analysis

Following the identification of ripples and cortical events, a window of 300 msec centered around each event peak was extracted for every event from the HPC, PFC and PPC. All ripple-centered epochs were subject to an artifact rejection stage. Epochs affected by artifacts were removed both manually through visual inspection and by implementing a statistical criteria in which epochs were rejected when their peak amplitude was higher than three scaled median absolute deviations (MAD) from the median peak amplitude value of the whole set of epochs (*43*). The total number of ripple and HFO events found varied considerably among conditions and is shown in Fig.1. Taking this into consideration and to avoid results being influenced by the different number of ripples and HFOs among conditions, both spectral analysis and granger analyses were performed using the same number of events (selected by the highest amplitude in PPC for HPC-PPC and PPC events, in HPC for HPC events) for every condition. Time-frequency analysis used the Short-time Fourier transform method to detect the changes of spectral power on HPC, PFC and PPC with respect to the time around the event. This was computed using the *ft_freqanalysis* function from the Matlab-based Fieldtrip toolbox (*44*) with a 100 msec Hanning window and a 90% overlap for a frequency range from 100 to 300 Hz with a 1Hz step. The resulting spectrograms were averaged among events per condition and the mean power value was extracted for a ± 50 msec window centered around the event peak from 100 to 250 Hz.

#### Spectral Granger Analysis

In order to determine the predictive power between the events corresponding to each brain region, the Spectral Granger Causality between them was computed for each condition. Granger Causality is a statistical hypothesis test for determining whether one time series is useful in forecasting another one for details please see (*45*). A 2.4 seconds window centered around each event peak was extracted for the simultaneous HPC, PFC and PPC signals. The same amount of events was used among conditions and they were ranked by the highest amplitude as described in the spectral analysis. A parametric Spectral Granger causality was computed by fitting a multivariate autoregressive model of order 10 selected based on the Akaike information criterion. This was performed with the *ft_mvaranalysis* function from Fieldtrip using the BSMART toolbox (*46*). The Fourier transform of the autoregressive model and its Granger causality were computed with the Fieldtrip functions *ft_freqanalysis* and *ft_connectivityanalysis* respectively. The resulting granger causality values for the 6 directionality combinations were split into frequency bands of 0-20Hz and 20-300Hz respectively. For each band the average granger value was calculated and reported. A non-parametric estimation of Spectral Granger causality based on (*47*) was also computed and shown in Fig. S4. This was done with a Multitaper method with Slepian tapers and a 2 Hz smoothing using the same Fieldtrip functions mentioned above.

#### Sharp wave ripple disruption

Ripples were detected online by bandpass-filtering the LFP signal from the CA1 pyramidal layer between 100 and 300 Hz. A user-defined threshold was applied to the rectified bandpass signal, to trigger stimulation. In the CON-D condition, bipolar stimulation was applied 200 ms to the ipsilateral or contralateral ventral commissure (AP -1.3, ML+/- 1.0, DV -3.8) after the ripple was detected. Because SWR-disruption tends to increase the number of SWR events (*33, 48*), one third of ripples were followed by two stimulation pulses, at 200 ms and 400 ms. This procedure was designed to approximately equalize the number of stimulations delivered across SWR-D and CON-D conditions. The closed-loop protocol was implemented in Cython (C-compiled Python-like language) introduced in the Open-Ephys signal chain by way of a Python Plugin (written by FP Battaglia, code available at https://github.com/MemDynLab/PythonPlugin).

## Supplement materials

**Fig. S1:**
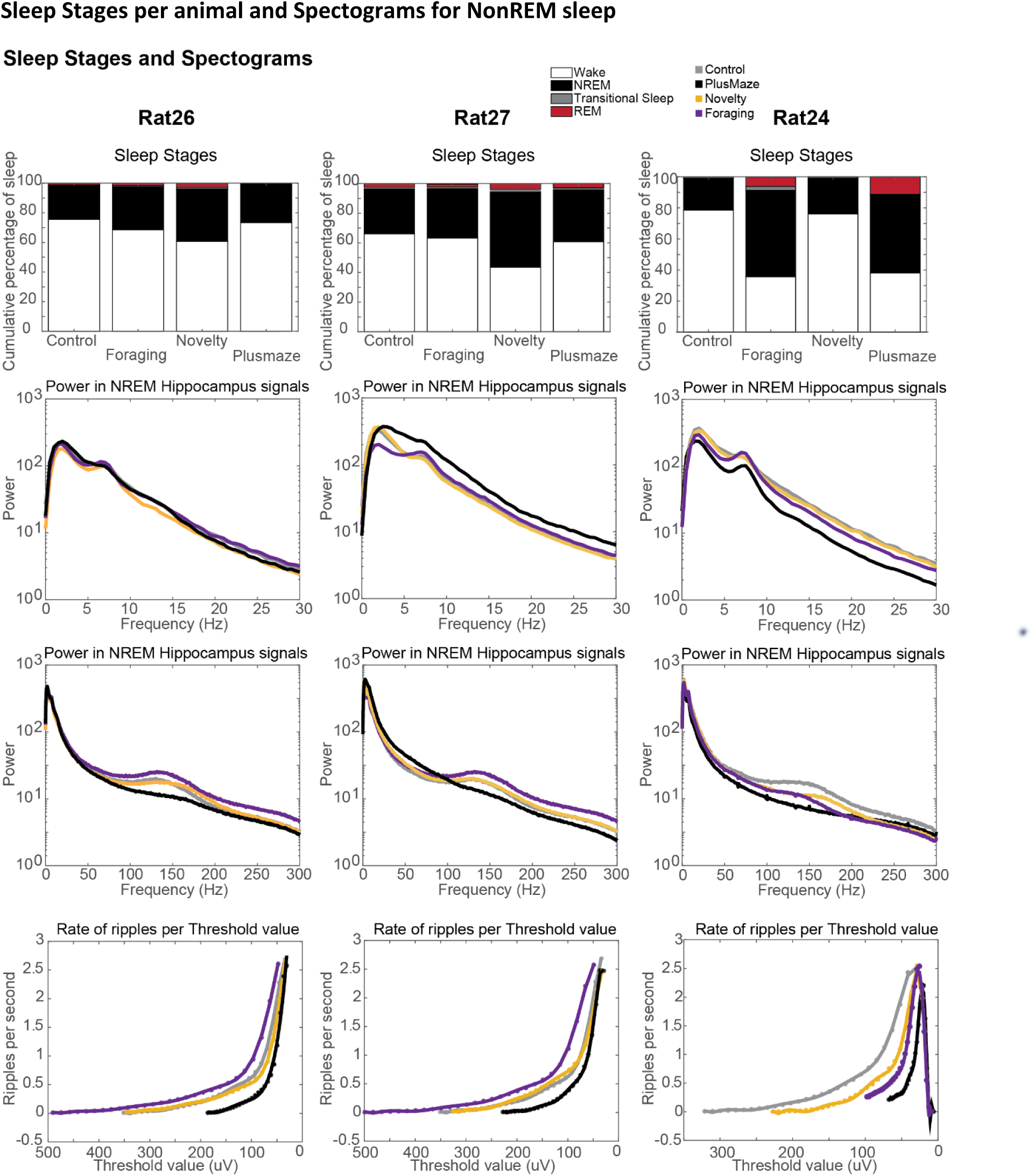
Shown are the (1) sleep stages and NonREM power spectra of (2) 0-30Hz as well as (3) 0-300Hz. And (4) number of detected ripples per threshold. All for each of the four conditions.

**Fig. S2:**
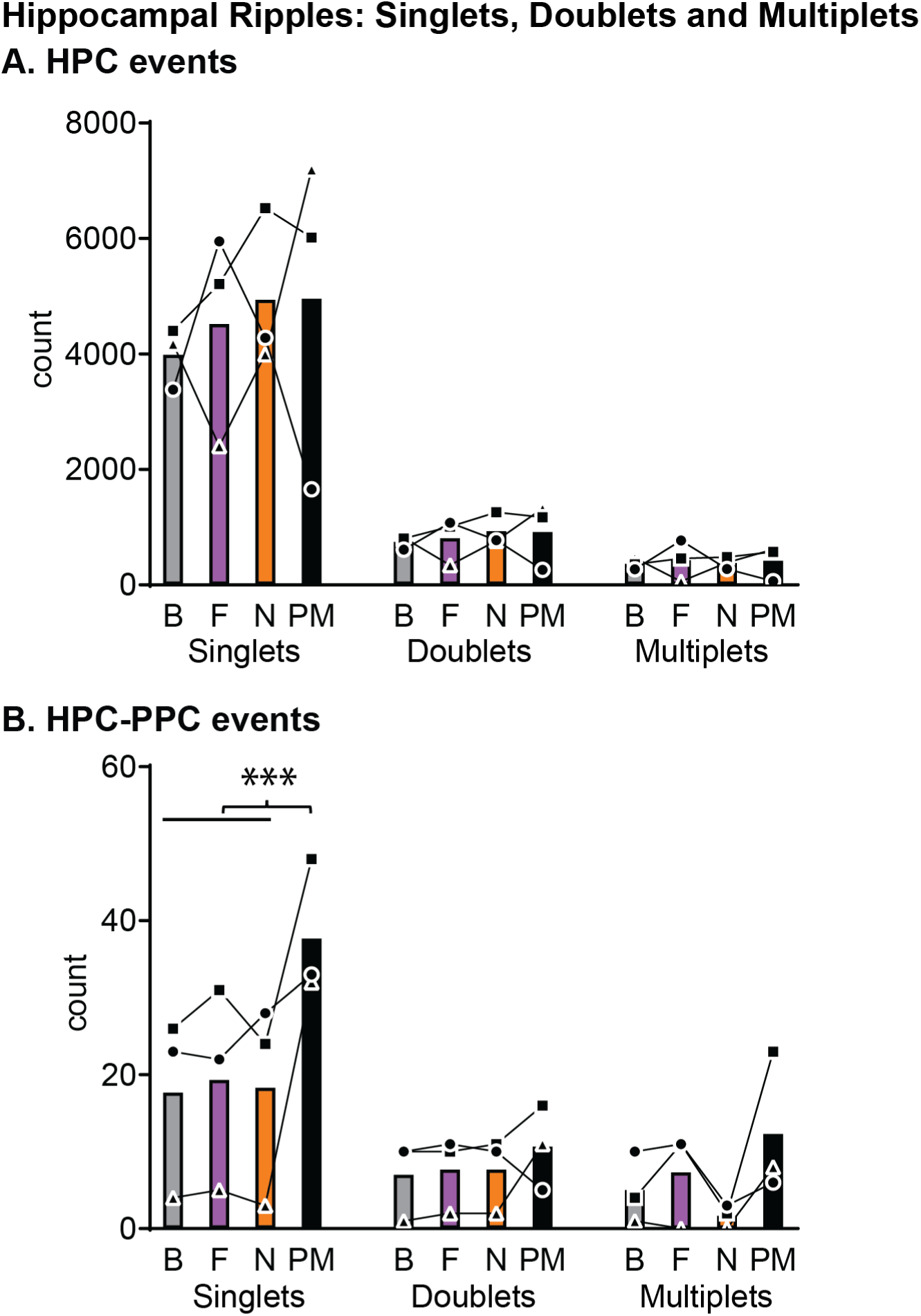
Here we classified the hippocampal ripples into singlets, doublets and multiplets (triplets and more) and then looked at HPC and HPC-PPC events across conditions. **A**. for HPC events there was an effect with more singlets than doublets or multiplets but no effect of condition. Types p=0.001 F_2,4_=84.69, other p> 0.9, F<0.2. **B**. for HPC-PPC events there was and type X cond interaction p=0.005 F_6,12_=5.62 as well as a type effect p=0.014 F_2,4_=15.18 and marginal condition effect p=0.063 F_3,6_=4.24. Only singlets showed a condition effect p=0.013 F_3,6_=8.88, **Orthogonal comparisons** ***p<0.001

**Fig. S3:**
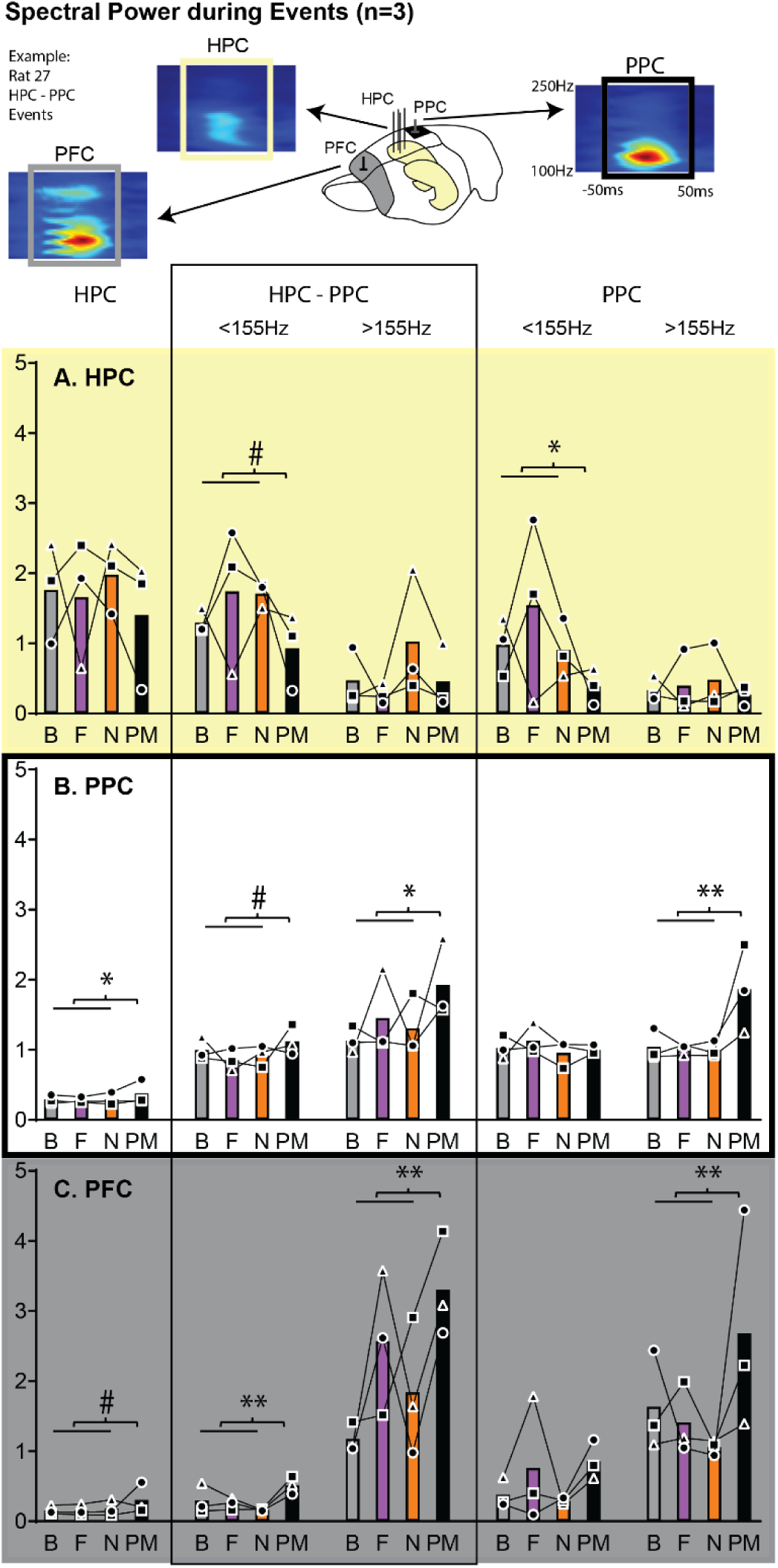
Spectral Power During Events split for single/coocurring: For all three brain areas normalized power. (for each animal across event type and brain area) for **100-250Hz ±50ms around the events** of the largest events in each condition (same number of events across conditions for each type). Shown for **A**. Hippocampus (HPC) **B**. posterior parietal cortex (PPC) and **C**. prefrontal cortex (AP 3.5, ML 0.5, ECoG, PFC). Fast PPC HF (independent if single or cooccurring) showed larger PFC and PPC but marginal smaller HPC spectral power as well as an increase in power after Plusmaze in PPC and PFC. Slow PPC HFO showed decreased HPC power after Plusmaze learning. **Full Model (BA, Type 5 levels, Cond)**: type p=0.019 F_4,8_=5.64, cond p=0.016 F_3,6_=7.93, type X BA interaction p<0.001 F_8,16_=22.58, type X cond interaction p=0.045 F_12,24_=2.24 (all other p>0.19). **Just including HPC-PPC and PPC events**: slow/fast p=0.002 F_1,2_=522.29, BA p=0.007 F_2,4_=21.36, cond p=0.004 F_3,6_=13.92, slow/fast X BA interaction p=0.003 F_2,4_=35.75, slow/fast X cond interaction p=0.075 F_3,6_=3.87; **for each brain area HPC-PPC and PPC events: HPC:** slow/fast p=0.065 F_1,2_=13.82, **PPC**: slow/fast p=0.024 F_1,2_=41.02, cond p=0.011 F_3,6_=9.38, **PFC:** slow/fast p=0.013 F_1,2_=75.48, cond p=0.047 F_3,6_=4.89 **Just including HPC-PPC events**: slow/fast p=0.075 F_1,2_=11.79, BA p=0.06 F_2,4_=6.19, slow/fast X BA interaction p=0.001 F_2,4_=56.26, BA X cond interaction p=0.073 F_6,12_=2.62, **Just including PPC events**: BA p=0.007 F_2,4_=22.15, cond p=0.032 F_3,6_=5.91, slow/fast X BA interaction p=0.026 F_2,4_=10.51, low/fast X cond interaction p=0.022 F_3,6_=6.96 **Orthogonal comparisons for condition**: #marginal p<0.1, *p<0.05, **p<0.01 Baseline (B), Foraging (F), Novelty (N), Plusmaze (PM).

**Fig. S4:**
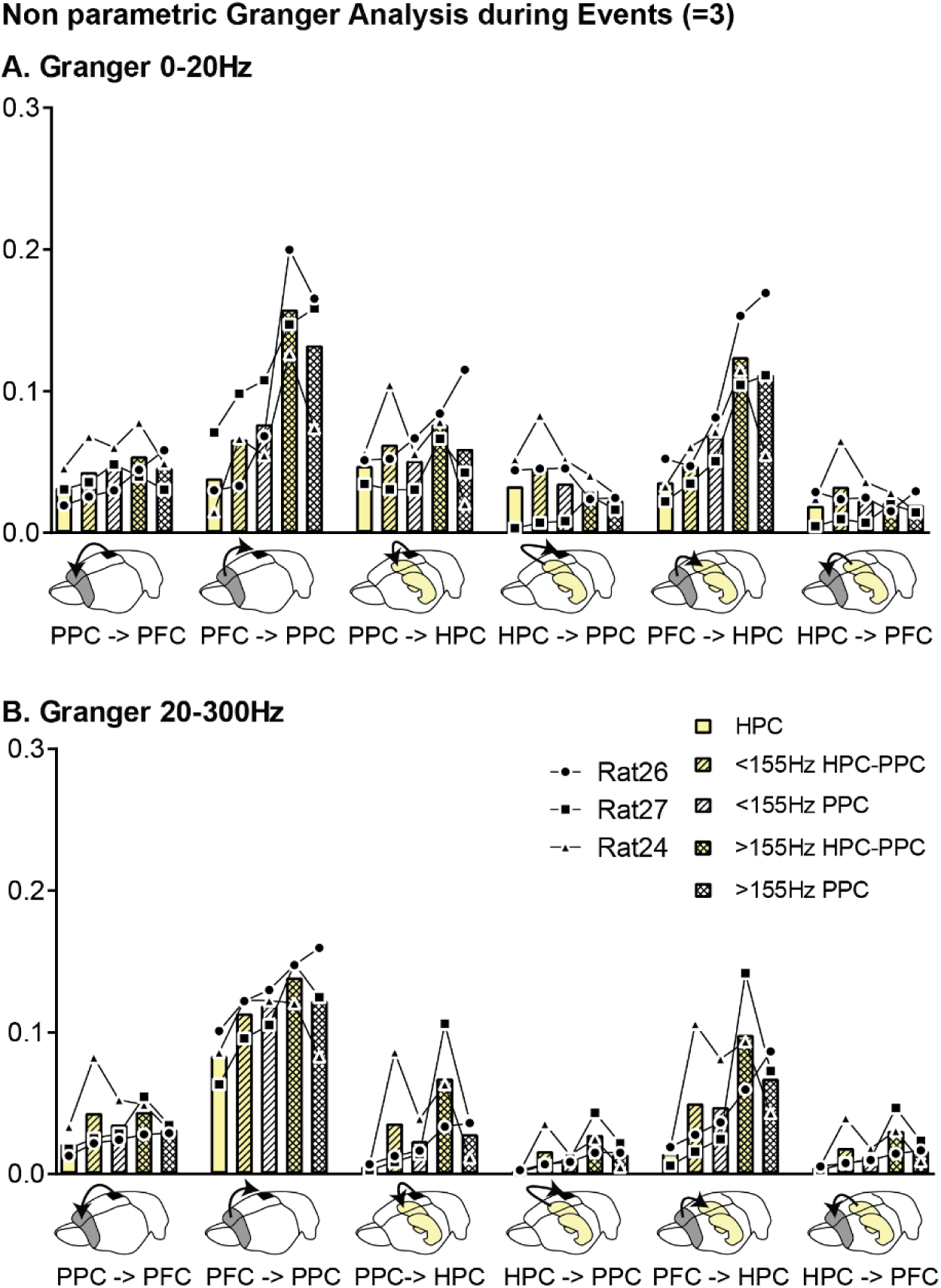
Show are the granger values across for 0-20Hz and 20-300Hz for the non-parametric analysis

**Fig. S5:**
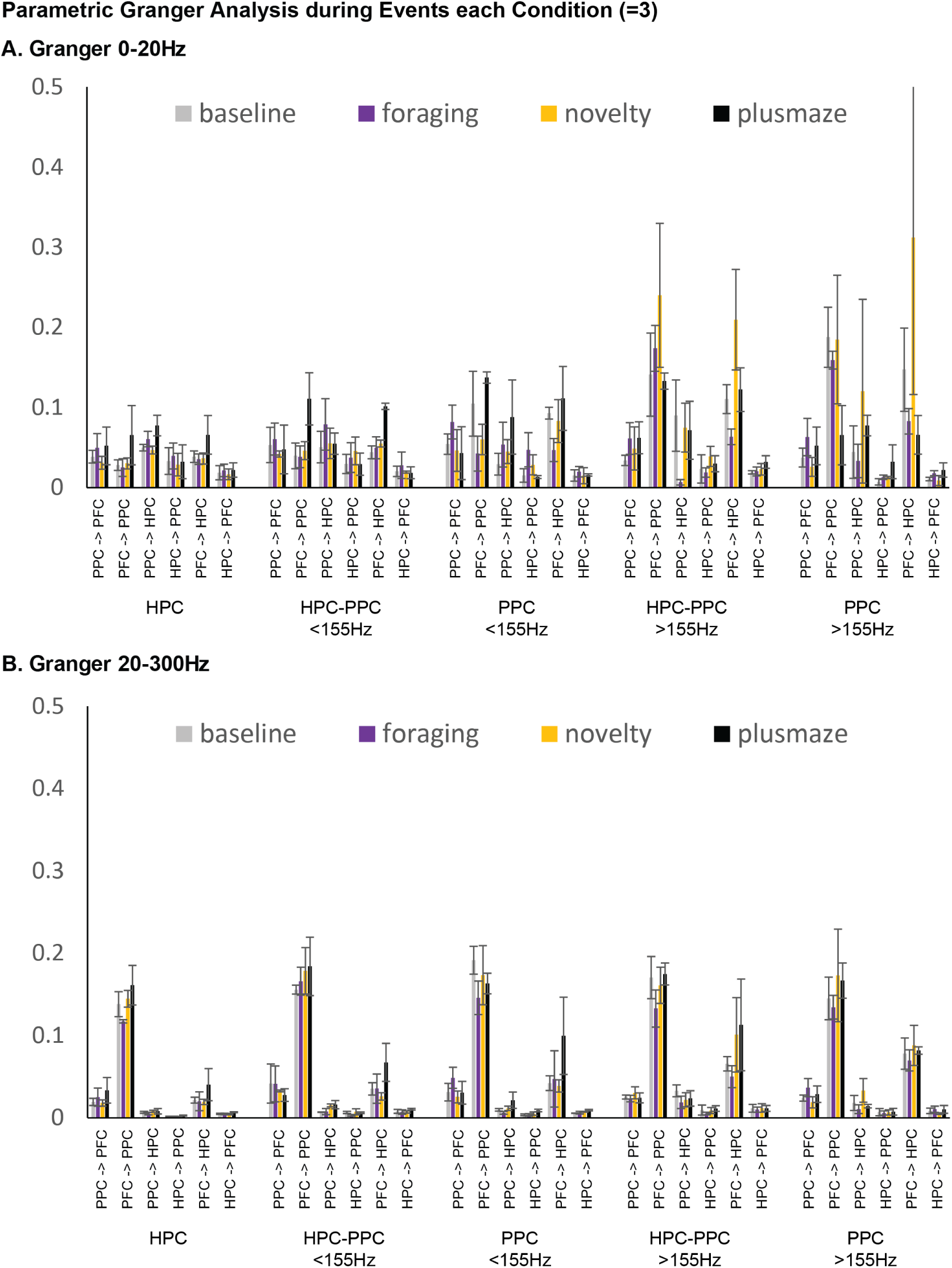
Show are the granger values across for 0-20Hz and 20-300Hz for the parametric analysis separately for each condition. Mean ± SEM

**Fig. S6:**
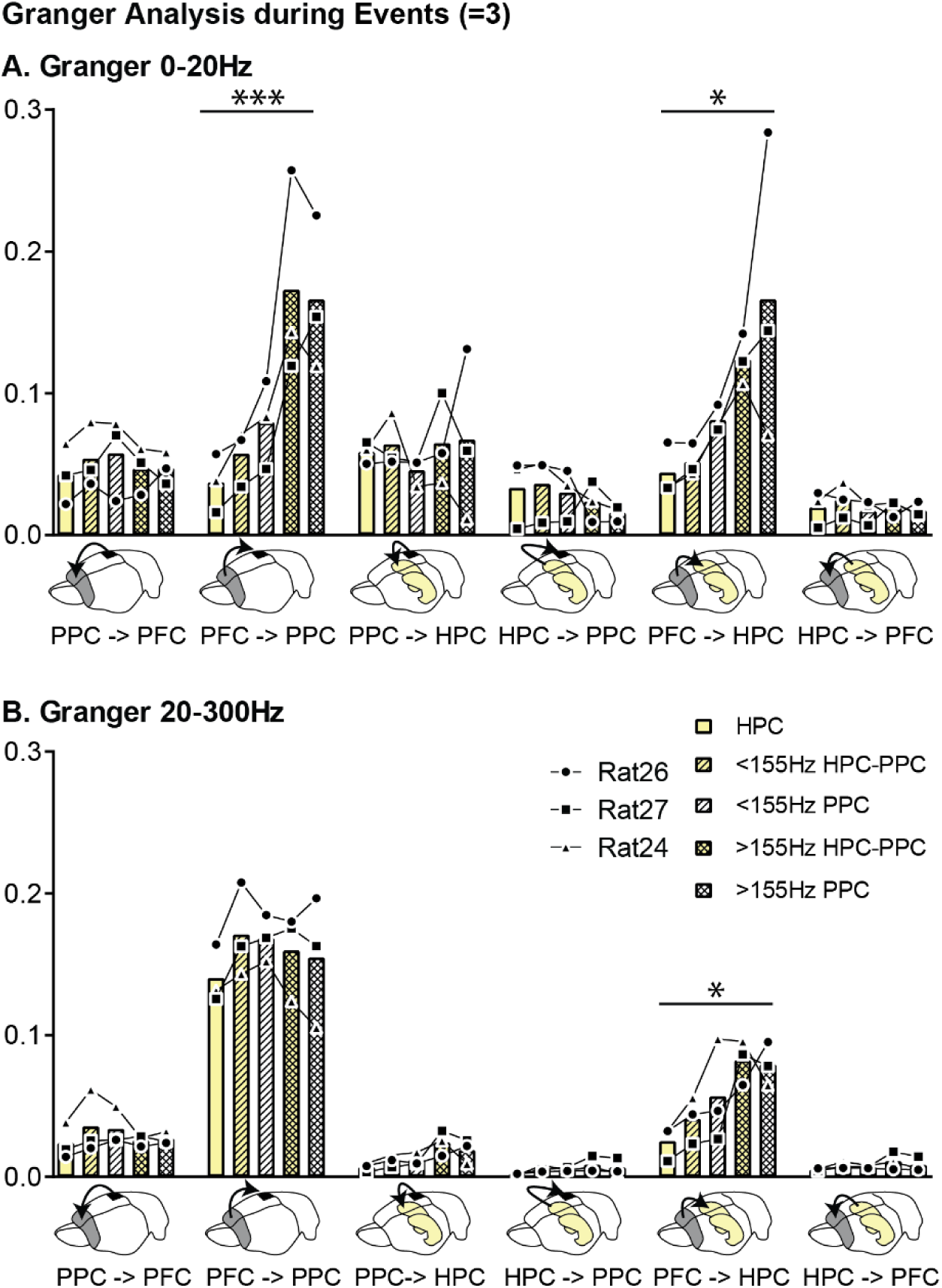
Granger Causality Analysis split for single/cooccurring. (parametric) is shown for both 0-20Hz and 20-300Hz oscillation bands of the events from Fig. 2 (single hippocampal ripples HPC [yellow], single posterior parietal cortex high frequency oscillations PPC [black and white shading] and cooccurring events HPC-PPC [yellow and black shading], divided for fast and slow PPC HFO) with the six possible directionalities. There was no difference in the conditions thus here an average per event type **A**. In the **slower frequencies** fast PPC HFO (both single and cooccurring) induced an increase in prefrontal cortex to hippocampal and to parietal. **B**. In contrast in the **faster frequency** band overall PFC to PPC was increased and PPC HFO (both single and cooccurring) showed an increase in PFC to HPC values **Full Model (Freq range, Directionality, Events)**: direct p<0.001 F_5,10_=26.06, events p=0.086 F_4,8_=3.02 direct X FR interaction p=0.001 F_5,10_=9.48, events X FR interaction p=0.072 F_4,8_=3.27, direct X events interaction p<0.001 F_20,40_=5.12, direct X event X FR interaction p<0.001 F_4,8_=4.90 For each oscillatory band separately: **0-20Hz** direct p=0.004 F_5,10_=7.52, events p=0.073 F_4,8_=3.25, direct X events interaction p<0.001 F_20,40_=5.61, **20-300Hz** direct p<0.001 F_5,10_=50.67, direct X events interaction p<0.001 F_20,40_=4.20, **Type effect**: *p<0.05,***p<0.001

**Fig. S7:**
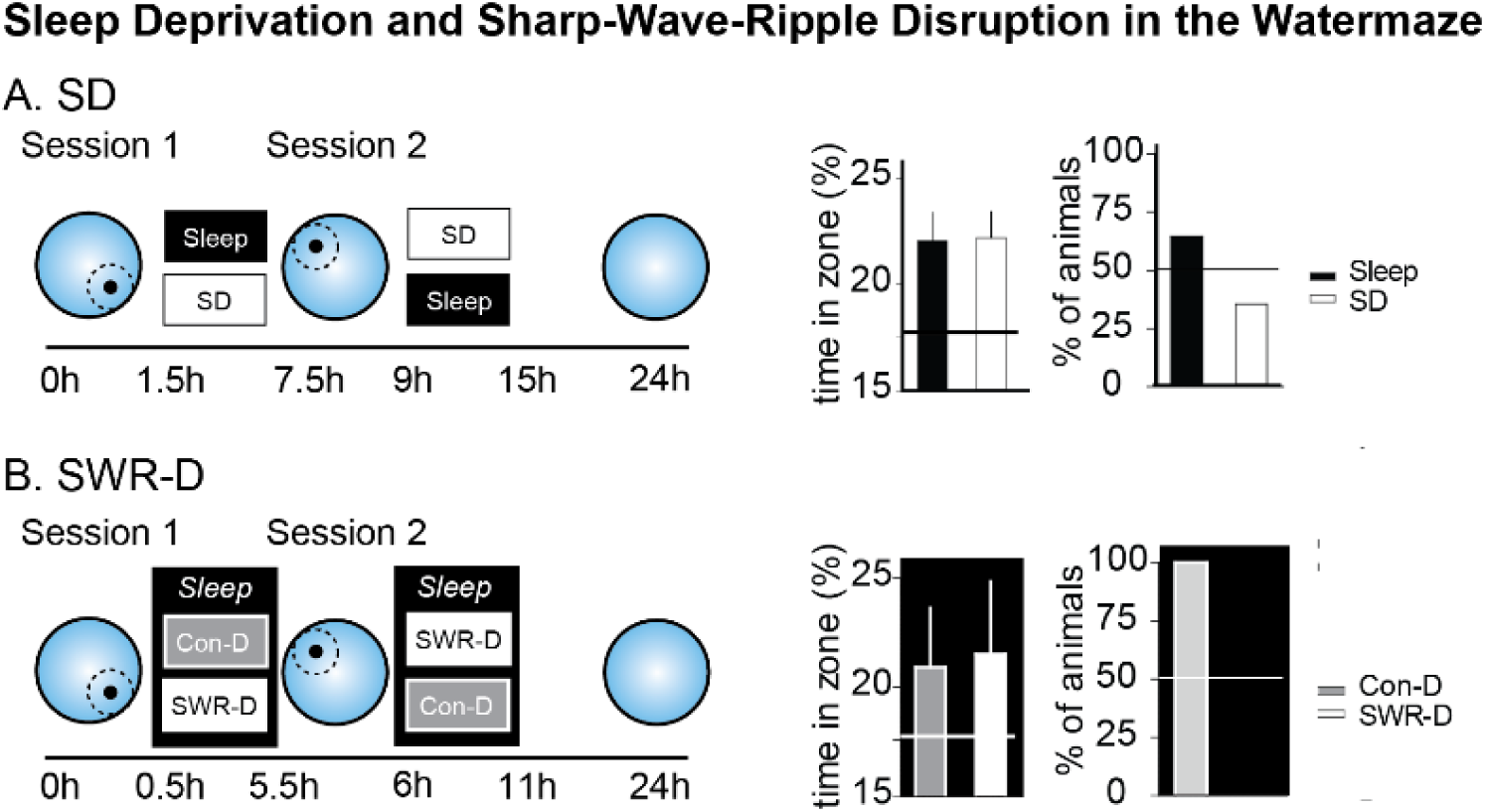
In addition to the Plusmaze experiments, we also ran the sleep deprivation and sharp-wave-ripple disruption (SWR-D) experiments in a watermaze paradigm. Unimplanted animals (n=32) were trained to one platform location (8 trials) after which they either were allowed to sleep or were sleep deprived (gentle handling) for 6h. Directly afterwards they were taught a second platform location (other side of the pool) and underwent the other condition (sleep/SD). Thus every animal was taught two platform locations, one of which was followed by sleep while the other was not. Data from (*1*), see also for more details on methods. Three implanted rats underwent the same procedure, but instead of sleep deprivation one location was followed by SWR-D and the other by our control disruption condition (Con-D). In both experiments animals spent the same amount of time at both platform locations, however both sleep deprivation and SWR-D led to the animals first swimming towards the other platform location (that was followed by uninterrupted sleep). This indicates that the watermaze memory while not dependent on sleep or sharp-wave-ripple activity in this paradigm, does change in quality or perhaps vividness due to sleep/SWR activity. Finally, these findings do replicate our main findings in the Plusmaze that SWR-D results in similar effects as sleep deprivation and thus may be the key mechanism of sleep’s effect on memory.

